# Identification of Immunodominant T-lymphocyte Epitope Peptides in HPV 1, 2 and 3 L1 Protein for Novel Cutaneous Wart Vaccine

**DOI:** 10.1101/2021.09.24.461620

**Authors:** Wu Han Toh, Chuang-Wei Wang, Wen-Hung Chung

## Abstract

**Background:** Common warts and flat warts are caused by the human papillomavirus (HPV). Peak incidence of wart infection occurs in schoolchildren aged 12-16, where prevalence can be as high as 20%. Traditional treatments aimed at destruction of wart tissue have low clearance rates and high recurrence rates. Occasional reports have even shown warts becoming malignant and progressing into verrucous carcinoma. Current licensed HPV vaccines largely target higher-risk oncogenic HPV types, but do not provide coverage of low-risk types associated with warts. To date, little attention has been given to the development of effective, anti-viral wart treatments.

**Objective:** This study aims to identify immunodominant T-lymphocyte epitopes from the L1 major capsid protein of HPV 1, 2 and 3, a foundational step in bioengineering a peptide-based vaccine for warts.

**Methods:** Cytotoxic T-cell and helper T-cell epitopes were predicted using an array of immunoinformatic tools against a reference panel of frequently observed MHC-I and MHC-II alleles. Predicted peptides were ranked based on IC_50_ and IFN-γ Inducer Scores, respectively, and top performing epitopes were synthesized and subjected to *in vitro* screening by IFN-γ enzyme-linked immunosorbent spot assay (ELISpot). Independent trials were conducted using PBMCs of healthy volunteers. Final chosen peptides were fused with flexible GS linkers in *silico* to design a novel polypeptide vaccine.

**Results:** Seven immunodominant peptides screened from 44 predicted peptides were included in the vaccine design, selected to elicit specific immune responses across MHC class I and class II, and across HPV types. Evaluation of the vaccine’s properties suggest that the vaccine is stable, non-allergenic, and provides near complete global population coverage (>99%). Solubility prediction and rare codon analysis indicate that the DNA sequence encoding the vaccine is suitable for high level expression in *Escherichia coli*.

**Conclusions:** In sum, this study demonstrates the potential and lays the framework for the development of a peptide-based vaccine against warts.

## 1. Introduction

Skin warts (verrucae) are benign lesions in the cutaneous epithelium. They are caused by the human papillomavirus (HPV), a small, non-enveloped, DNA virus that is easily transmitted through skin-to-skin contact [1]. Peak incidence of wart infection occurs in schoolchildren aged 12-16, where prevalence can be as high as 20% [2]. While two-thirds of warts regress spontaneously within a few years, they may cause significant morbidity in affected individuals. Occasional reports have even shown warts becoming malignant and progressing into verrucous carcinoma [3].

While there are more than 200 HPV genotypes, only a handful are responsible for cutaneous warts. Common warts are most frequently associated with HPV types 1, 2, and 7, plantar warts with types 1 and 2, and flat warts with types 3 and 10. Other higher-risk HPV types are oncogenic and are a principal cause of cervical cancer [4, 5]. Differentiation of HPV types is based on the nucleotide sequence of the L1 major protein [6]. The L1 protein, along with the L2 minor component, forms pentameric capsomer subunits that make up the capsid shell of the virus. L1 has been shown to spontaneously self-assemble into virus-like particles (VLPs) that can act as highly potent immunogens, presenting an exterior surface nearly indistinguishable from the native virion [7]. As such, the L1 protein has been widely used in existing VLP-based HPV vaccines and is the target for novel subunit vaccines.

Thus far, vaccine development efforts for HPV have been centered on higher-risk HPV types. There are currently 3 licensed VLP-based HPV vaccines: Cervarix® bivalent vaccine, and Gardasil® Quadrivalent and 9-valent vaccine [8].These vaccines all possess a prophylactic effect against HPV types 16 and 18, which together account for approximately 70% of cervical cancer cases [9]. Because VLPs lack the viral genome but have a morphologically similar outer structure to the virus, they are both noninfectious and highly immunogenic. Recent studies have also sought to develop effective therapeutic HPV vaccines, as well as vaccines that improve accessibility to developing countries [10–13]. In comparison, little attention has been afforded to vaccine development for low-risk HPV types. To date, no effective antiviral drugs are available for wart treatment.

Instead, existing treatment modalities are largely aimed at destruction of wart tissue. Cryotherapy by liquid nitrogen is the most common approach and involves inducing cold injury to warts [14]. Freezing treatment is repeated every 2-3 weeks until warts have cleared. Salicylic acid application is also commonly used to promote exfoliation of epidermal cells and is postulated to have an added mechanism of stimulating host immunity [15]. Studies have proven salicylic acid to be somewhat more effective than placebo [16]. Nonetheless, conventional treatments can be painful and inefficient, with low clearance rates and high recurrence rates [17]. Moreover, treatment has the potential to cause moderate to severe scarring, leading to severe deficits in cosmesis [3]. These risks have led many physicians to opt for a wait-and-see approach, leaving patients with cosmetic insecurity and physical discomfort. Others have turned to alternative techniques such as laser therapy and oral isotretinoin, but issues over recurrent infection remain [17–19]. Evidently, there is a need for a prophylactic, nondestructive wart treatment. We observe that the creation of a treatment with these properties can be achieved by adapting the vaccine design approaches currently employed for higher-risk HPV types.

This study aims to identify immunodominant T-lymphocyte epitopes from the L1 protein of HPV 1, 2 and 3, as the first step for bioengineering a peptide-based vaccine for warts. In recent years, peptides have become increasingly attractive vaccine candidates with the potential to overcome the disadvantages of traditional vaccines, including those based on VLPs. Compared to vaccines using eukaryotic expression systems, peptide-based vaccines are easily produced in *Escherichia coli*, and their greater thermal stability eliminates the need for cold chain [20]. The lack of redundant elements also minimizes unwanted allergic reactions and allows the vaccine to be tailored for more focused immune responses [21, 22]. Beyond these advantages, the inclusion of multiple epitopes allows for a single peptide vaccine to target several strains, which is crucial to maximizing coverage across wart-causing HPV types with low sequence similarity [23].

Epitope selection is the first and most crucial stage in designing peptide-based vaccines. Effective epitopes must be able to induce high levels of humoral and cellular immune responses. In this study, we employ a peptide screening method relying on immunoinformatic tools that predict MHC-I and MHC-II binding affinity, followed by *in-vitro* testing by enzyme-linked immunosorbent spot (ELISpot) assay [24]. This two-step process reduces the number of experiments needed, thereby facilitating efficient candidate epitope identification. Following *in-silico* screening, ELISpot assay was performed to evaluate the response of human CD4^+^ and CD8^+^ T-lymphocytes to the peptide stimuli. The assay provides quantitative measurements of the secretion of IFN-γ, an abundant cytokine produced by Th1 cell with a critical role in activation of innate and adaptive immunity. IFN-γ ELISpot assays are used extensively for screening of immune responses in vaccine development and have been shown to be both sensitive and reproducible [25, 26].

## 2. Materials and Methods

### 2.1 Sequence Retrieval

The amino acid sequences of the L1 protein for HPV 1a, 2, and 3 were retrieved from the National Center for Biotechnology Information database (http://www.ncbi.nlm.nih.gov/) with accession numbers of CAA01195.1, ABO14919.1, and P36731.1, respectively.

### 2.2 T-cell Epitope Prediction

T-cell epitope prediction was performed separately for each of the retrieved sequences, due to low sequence similarity among the HPV types. The prediction process is summarized by the flowchart depicted in Figure 1.

**Fig 1.**
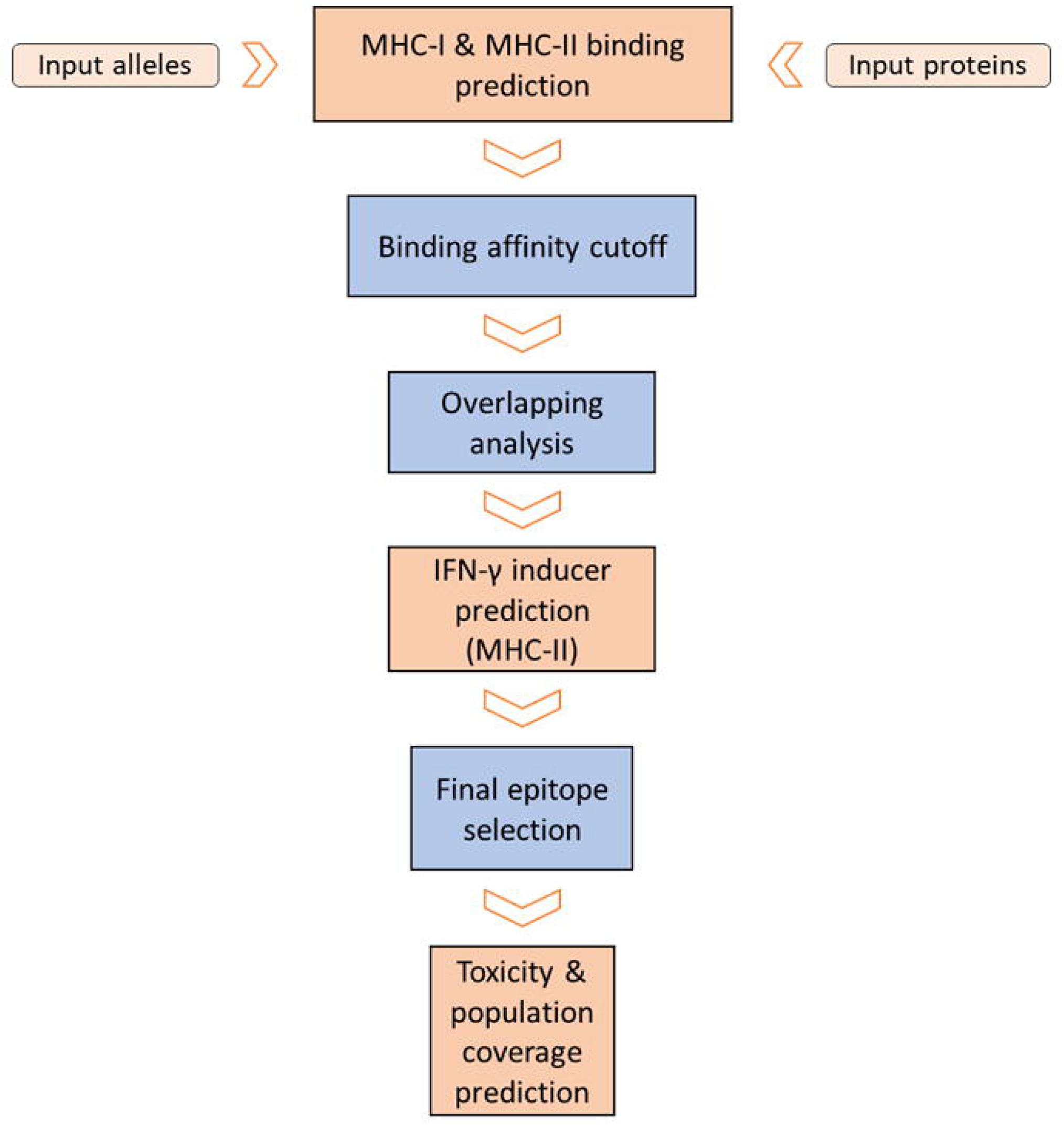
Flowchart for bioinformatics analysis in the study. The process was repeated to predict epitopes for HPV 1, 2, and 3.

#### 2.2.1 MHC Class I Epitope Prediction

MHC-I binding epitope prediction was performed using three methods from the IEDB (www.iedb.org) database (Table 1). Predictions of 8-11 mer epitope binding sites were made against a reference panel of 27 frequently observed MHC-I HLA alleles, with a reported population coverage greater than 97% [27]. Predicted epitopes were ranked according to their *half maximal inhibitory concentration* (IC_50_) scores, and peptides below the MHC-I allele specific affinity cutoff thresholds were excluded from further analysis [28, 29]. From this filtered list, overlapping sequences from the three prediction methods were selected.

**Table 1.** Methods used for T-cell epitope binding prediction.

|  | Server | Link | Prediction Method | References |
| --- | --- | --- | --- | --- |
| **MHC-I Binding** | IEDB | http://tools.iedb.org/mhci/ | NetMHC (ANN) 4.0 – ANNs | [31-36] |
|  |  |  | NetMHCI Pan BA 4.0 – ANNS | [37-41] |
|  |  |  | PickPocket – binding specificities based on receptor pocket similarities | [42] |
| **MHC-II Binding** | IEDB | http://tools.iedb.org/mhcii/ | IEDB Recommended 2.22 – consensus approach combining ANN, SMM, combinatorial library, and pocket profiles | [43-49] |
|  | NetMHCIIpan | http://www.cbs.dtu.dk/services/NetMHCIIpan-3.2/ | NetMHCIIpan 3.2 - ANNs | [43] |

#### 2.2.2 MHC Class II Epitope Prediction

Two servers, IEDB and NetMHCIIpan 3.2, were used for prediction of 15 mer MHC-II binding epitopes (Table 1). A similar reference panel of 27 MHC-II HLA alleles with a population coverage greater than 99% was used for analysis [30]. Predicted peptides with an adjusted percentile rank greater than 10 were filtered out, and subsequent overlapping analysis of the results of the two prediction methods was performed.

#### 2.2.3 Evaluation of Selected Epitopes

Overlapping MHC-I and MHC-II epitopes were reranked based on mean IC_50_ and IFN-γ Inducer Score, respectively. The potential of MHC-II epitopes to induce IFN-γ production was evaluated with the IFNepitope server (https://crdd.osdd.net/raghava/ifnepitope), using a Support Vector Machines (SVMs) based method [50]. This study recognizes the importance of CD4^+^ cell activation in vaccine formulation. Accordingly, 15 MHC-II binding peptides with the highest IFN-γ Inducer Score, and 5 MHC-I binding peptides with the lowest mean IC_50_ were chosen for both HPV type 1 and 2. Only 5 peptides with the highest binding affinities were selected from HPV type 3, in accordance with the strain’s relatively lower prevalence of infection. Lastly, the toxicity (http://crdd.osdd.net/raghava/toxinpred/multi_submit.php) of the chosen peptides were assessed, and peptides with a positive toxicity prediction were excluded [51]. In total, 45 peptides were elected for *in vitro* testing.

### 2.3 Peptide Synthesis

Peptide sequences were synthesized by Kelowna International Scientific (Taiwan) with more than 90% purity. Peptides were received in lyophilized form, and stock solutions were created by dissolving 40 mg/mL in DMSO. Peptides were stored at −20 °C until use, where they were diluted to 1 mg/mL in PBS.

### 2.4 In vitro Immunodominant Peptides Screening

To screen the candidate peptides, IFN-γ ELISpot assay (Mabtech) was performed using PBMCs of healthy human volunteers, in 4 independent trials. Isolated T-cells were seeded 2 × 10^5^ in 200 μL medium (RPMI, 5% AB serum, 1% PSA) with IL-2 and IL-7 cytokines in round bottom 96 well plates. 4 μL of peptide solution or PBS control was added to the wells, so that each peptide was present in 2 wells. Dendritic cells (DC) in 15 mL medium were seeded in a T75 cell culture flask. Cultured cells were incubated at 37 °C. Cell medium was changed and peptides were added every 2 days. After 7 days, following DC maturation, T-cells and DCs were cocultured in a 10:1 ratio. Cells were cultured for another 6 days.

ELISpot PVDF plates (MSIPS4510, Millipore) were activated with 30 μL 35% EtOH for 1 minute. Following washing, 100 μL of 15 μg/mL 1-D1K coating antibody was added to each well and allowed to react overnight. After blocking with 200 μL medium with 10% AB serum for 30 min, cells were seeded 1 x 10^5^ per well and stimulated with 8 μL peptide, 2 μL OKT-3, or 8 μL medium. Spots were detected using a biotinylated antibody (7-B6-1-biotin) against streptavidin conjugated ALP and visualized by the addition of a colorimetric substate (BCIP/NBT).

Plates were scanned and the number of spot forming cells (SFCs) in each well were counted by an ImmunoSpot reader (CTL Cellular Technology). The stimulation index of each peptide was calculated as the ratio of SFCs in peptide-stimulated cells to that of non-stimulated cells. Peptides with an SI value greater than 2 were regarded as positive and immunodominant.

### 2.5 Design of Recombinant Polypeptide Vaccine and In Silico Cloning

Of the peptides that passed ELISpot screening, 7 were included in the design of a polypeptide HPV 1, 2, and 3 vaccine. Peptides were fused with a flexible glycine-serine linker (GS), as studies have reported they provide the best structure and stability for fusion proteins [52]. The physiochemical properties, including stability, molecular weight, and isoelectric point, of the resulting polypeptide vaccine were evaluated using ProtParam server (https://web.expasy.org/protparam/), and the vaccine’s global population coverage was analyzed using the IEDB tool (http://tools.iedb.org/population/) [51, 53]. Protein solubility upon expression in *E. Coli* was predicted using the ccSOL omics server (http://s.tartaglialab.com/page/ccsol_group), with a reported 74% accuracy [54]. Finally, AllergenFP (http://ddg-pharmfac.net/AllergenFP/) was used to ensure the vaccine had no allergenic properties [55].

To prepare for expression in *E. coli*, the polypeptide sequence was reverse translated. A hexahistidine tag (6x-His) was attached upstream of the vaccine sequence for ease of purification, and the entire sequence was codon optimized through the EMBOSS backtranseq tool (https://www.ebi.ac.uk/Tools/st/emboss_backtranseq/) [56]. To verify the potential for high level protein expression, the optimized DNA sequence was evaluated for codon adaptation index (CAI), GC content, and codon frequency and distribution (CFD) using the GenScript Rare Codon Analysis Tool (https://www.genscript.com/tools/rare-codon-analysis) [13]. Finally, restriction sites of EcoRI and XbaI were inserted at the start of the sequence, while SpeI and PstI were inserted at the end.

## 3. Results

### 3.1 T-cell Epitope Prediction

In the first phase of this study, a variety of bioinformatics tools were employed to predict and filter T-cell epitopes from the L1 protein of wart-causing HPV strains (Table 2). During this prediction phase, an emphasis was placed on binding affinity, as it is both an important factor for vaccine construction and can be reliably predicted by existing tools. MHC-I and MHC-II binding predictions were consolidated from multiple methods to ensure accurate predictions. Of the 45 highest scoring candidate peptides, 44 were successfully synthesized and subjected to *in vitro* screening.

**Table 2.**
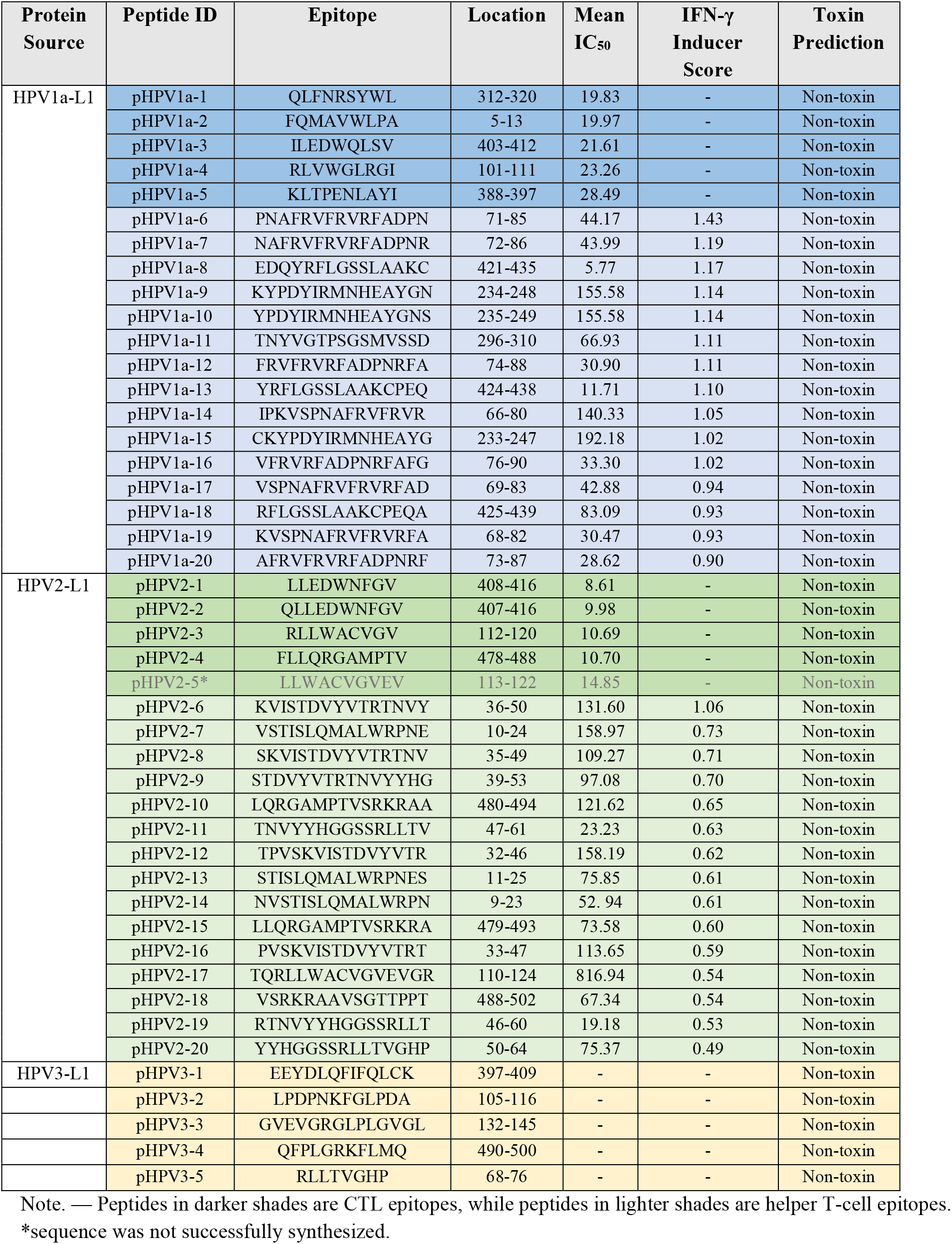
Predicted epitopes from HPV 1, 2, and 3 L1 major capsid proteins.

### 3.2 In vitro Immunodominant Peptides Screening

Elected peptides were subjected to screening by IFN-γ ELISpot assay, where cells were stimulated with PBS, peptides, or OKT-3 positive control. Results from independent trials of 4 human volunteers were averaged. Of the 44 peptides screened, 11 produced an SI value greater than 2 and were therefore considered immunodominant (Fig. 2). The distribution of targets for the immunodominant peptides were 1 for MHC-I HPV 1a, 7 for MHC-II HPV 1a, 2 for MHC-II HPV 2, and 1 for HPV 3.

**Fig. 2.**
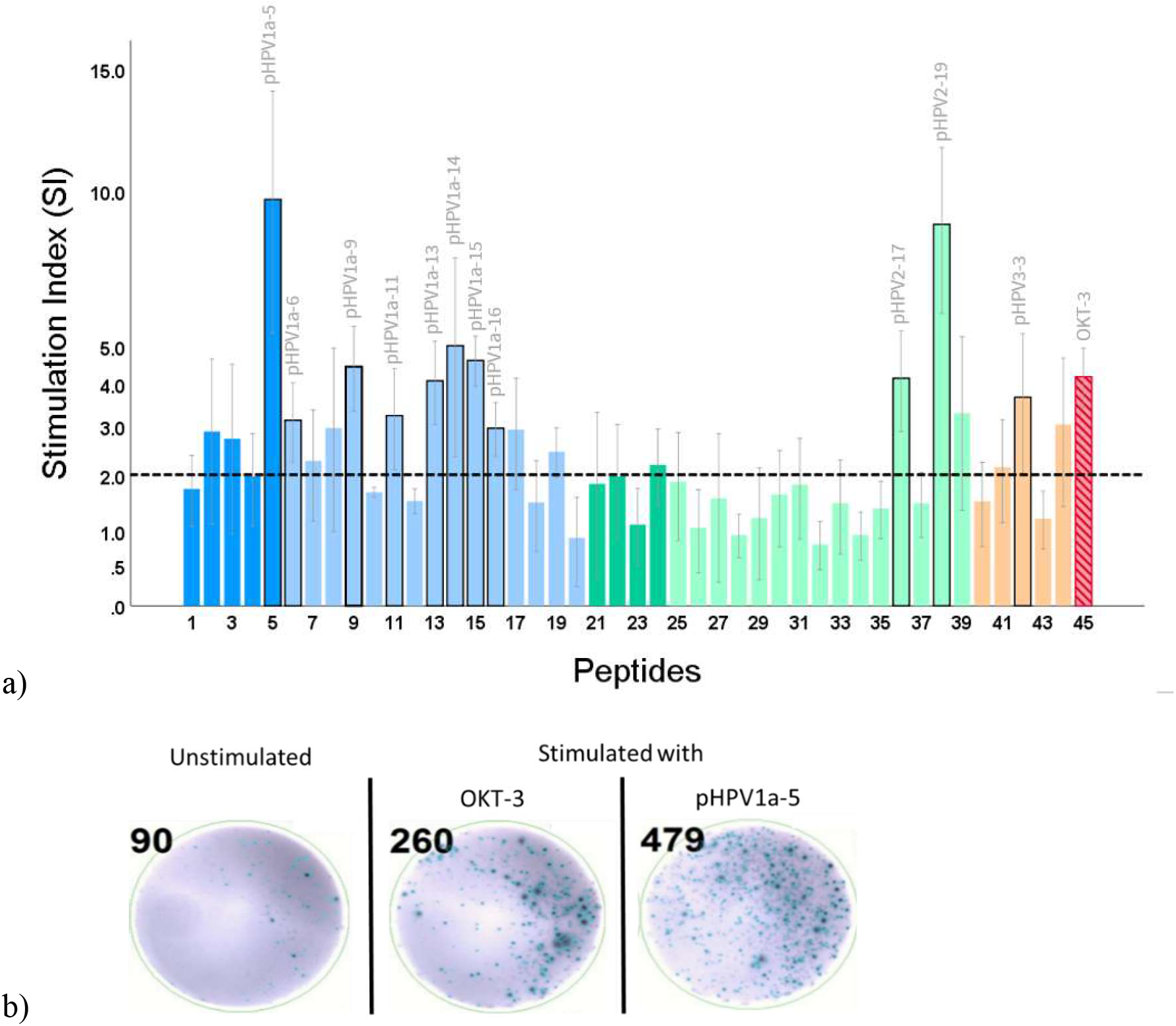
Immunodominant peptides screening via ELISpot assay. a) PBMCs obtained from healthy human volunteers were stimulated with peptides selected from the L1 protein of HPV 1a (blue), HPV 2 (green), and HPV 3 (orange). The stimulation index was calculated as the ratio of SFCs in peptide stimulated cells and PBS mediums-stimulated cells. The threshold for immunodominance was set at SI > 2, as labeled in the figure. b) A representative image of wells analyzed by a CTL ImmunoSpot reader is shown, illustrating the number of spots observed in a non-stimulated well, an OKT-3 stimulated well, and a well stimulated by an immunodominant peptide, pHPV1a-5.

### 3.3 Sequence Selection for Vaccine Design

In preparation for vaccine design, further screening of the 7 MHC-II epitopes for HPV 1a was performed to reduce redundancy. Using the IEDB population coverage prediction tool (http://tools.iedb.org/population/), it was shown that peptides pHPV1a-6 and pHPV1a-15 had global population coverages of 96.90% and 77.19%, respectively, significantly higher than the other 5 peptide sequences. Thus, they were chosen for the vaccine. Next, the lack of immunodominant MHC-I epitopes targeting HPV 2 was resolved by the inclusion of pHPV2-4 in the vaccine. While the mean SI of pHPV2-4 fell slightly below the threshold, the peptide yielded positive results in 2 of the 4 subjects. Ultimately, 7 peptides were included in the final vaccine design (Table 3).

**Table 3.**
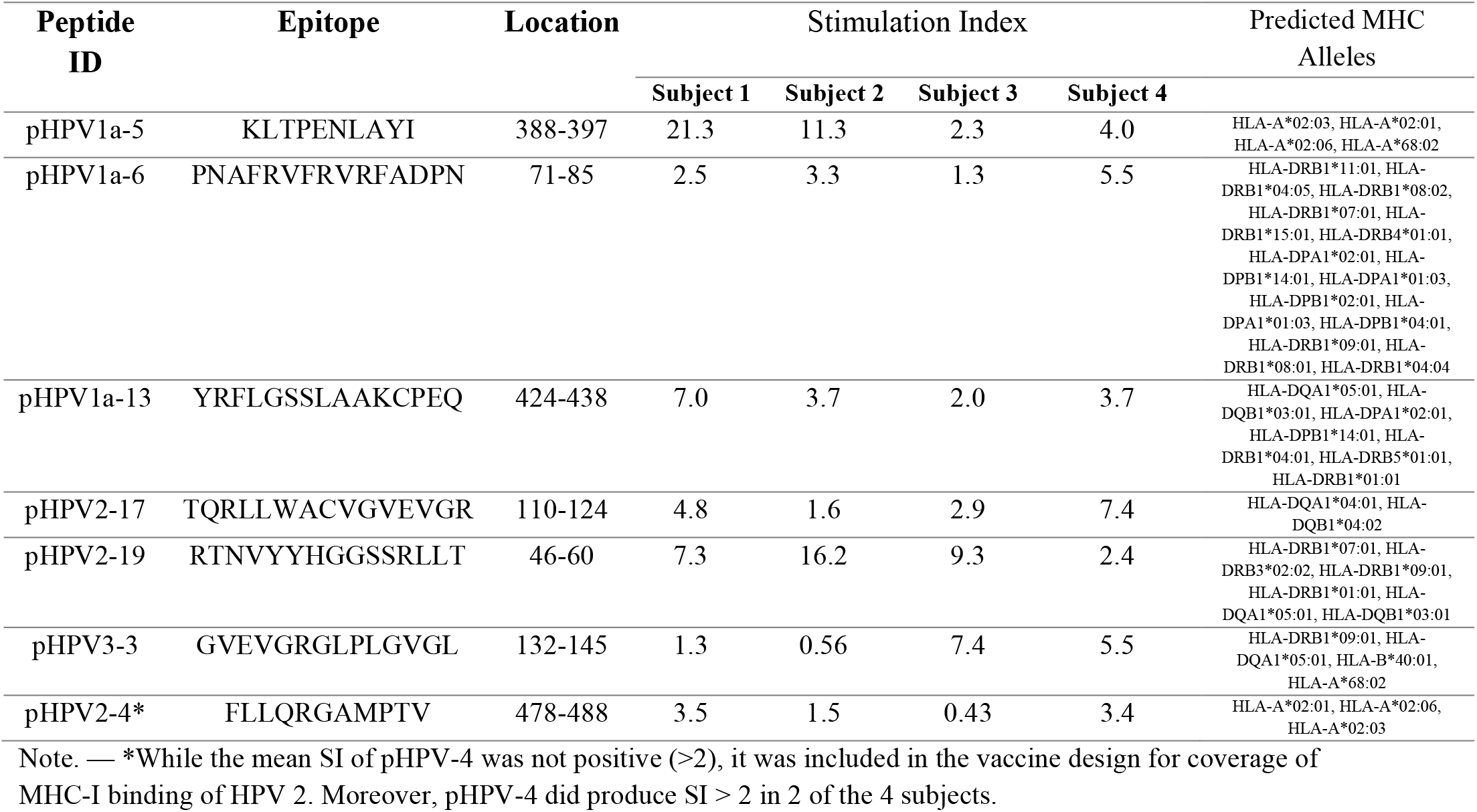
Peptides selected for vaccine design.

### 3.4 Design of Recombinant Polypeptide Vaccine and In Silico Cloning

Figure 3 illustrates the construct of the candidate polypeptide vaccine. Inclusion of multiple peptides allows for simultaneous coverage of HPV types 1, 2, and 3. Results from server analysis of the vaccine’s physiochemical properties show that it has a molecular weight of 12.04 kDa, a theoretical pI of 10.27, and an instability index of 39.70, indicating that the vaccine is stable (Table 4). Furthermore, the vaccine is also predicted to have high global population coverage (99.63%), and decent solubility propensity (54%). The vaccine is classified as a nonallergen.

**Fig. 3.**
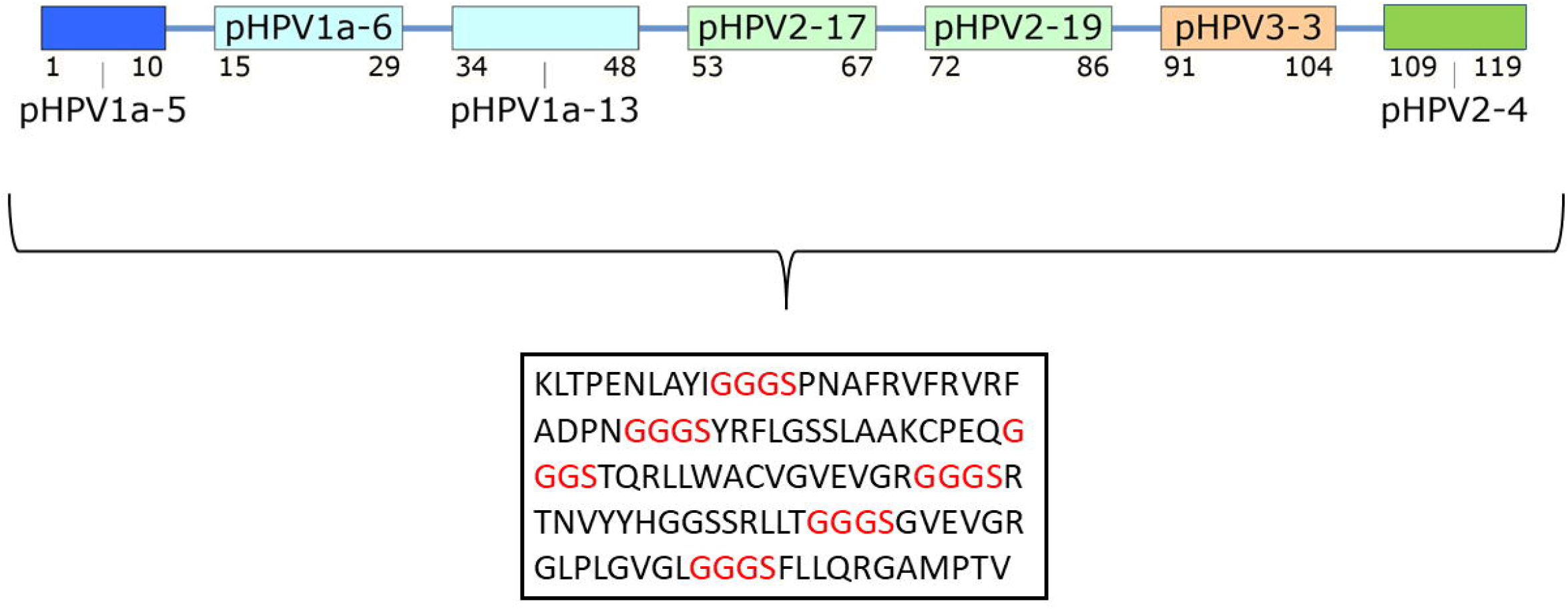
Structure and sequence of recombinant peptide vaccine. Chosen peptides were joined in series with flexible GS linkers, as labelled in red. The total polypeptide is 119 amino acids long.

**Table 4.** Physiochemical properties, population coverage, and allergenicity of recombinant vaccine.

| *Vaccine Properties* | *Value* |
| --- | --- |
| *Molecular weight (Da)* | 12044.63 |
| *No. of amino acids* | 119 |
| *Instability index* | 39.70 (Stable) |
| *Gravy* | -0.062 |
| *Aliphatic index* | 76.13 |
| *Theoretical pI* | 10.27 |
| *Solubility propensity* | 54% |
| *Global population coverage* | 99.63% |
| *Allergenicity* | Probable Non-Allergen |

Following reverse translation, the codon optimized gene for the vaccine was evaluated for codon adaptation index, GC content, and codon frequency distribution with *E. Coli* selected as the expression host (Fig. 4). Results from these parameters indicate the sequence is suitable for cloning and expression in *E. coli*.

**Fig. 4.**
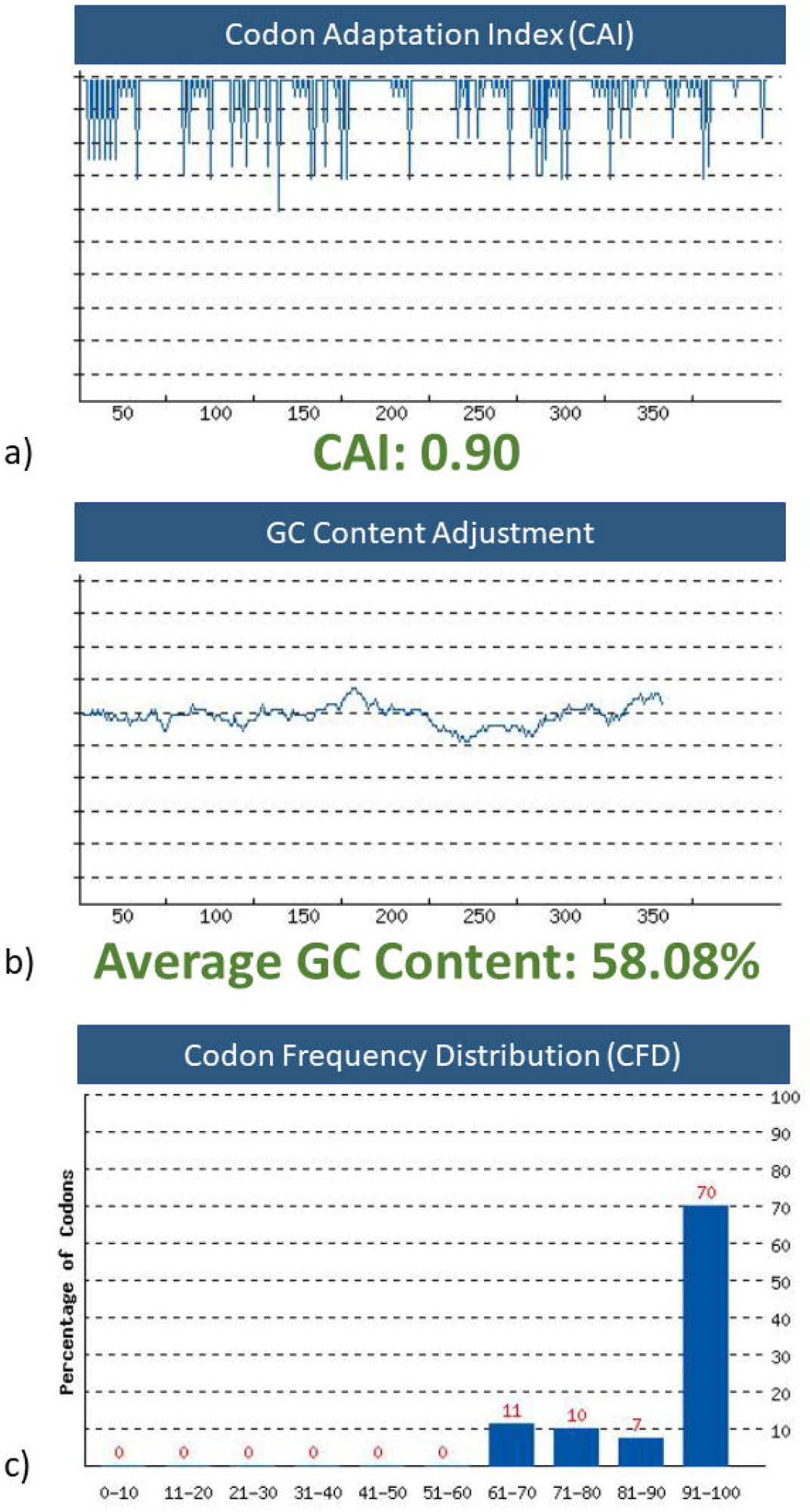
Evaluation of parameters for high-level protein expression in *E. coli*. a) The sequence has a CAI value of 0.90. CAI is the distribution of codon usage frequency along the length of the coding region, and a value > 0.8 is considered good for expression. b) The average GC content of the sequence is 58.08%, and there are no peaks outside of the ideal range of 30 – 70%. c) The usage of codons with frequency scores lower than 30 are likely to adversely affect expression efficiency. The percentage of low frequency codons in this sequence is 0%.

## 4. Discussion

Existing wart treatments have low clearance rates and high recurrence rates, and as such, are increasingly seen as non-viable options for recalcitrant infections [17]. Licensed VLP-based HPV vaccines on the market target high-risk oncogenic HPV types, and while studies have reported limited positive responses against anogenital and cutaneous warts, their efficacy is hindered by a lack of specific coverage for verrucae-causing types, namely, HPV types 1, 2, and 3 [57, 58].

In this study, we aim to repurpose the peptide-based vaccine design methods currently employed for high-risk HPV types to develop a novel, cutaneous wart vaccine. The value of such an approach is derived from its therapeutic and prophylactic effects, its non-destructive nature, as well as its ease of production and transport compared to traditional vaccines [20]. Furthermore, by eliminating the need for a strict treatment regimen, the approach holds the potential to relieve patients and physicians from the frustrations caused by problematic warts [59].

The proposed recombinant polypeptide vaccine is the product of a peptide screening process combining both immunoinformatics and *in vitro* assays. In recent years, the fields of genomics and bioinformatics have evolved to enable computational screening of epitopes considering a diversity of HLAs [60]. The reliability of such methods can be increased by analyzing the results from multiple prediction tools. This study uses immunoinformatic tools to narrow the epitope pool to a manageable number for *in vitro* tests, to facilitate an efficient and accurate experimental design.

Using this process, the final vaccine design consists of 7 T-lymphocyte epitopes, including both CTL and T-helper cell epitopes, selected for their immunodominance via human IFN-γ ELISpot assays. The resultant vaccine is multi-target, with epitopes designed to elicit specific immune responses to HPV types 1, 2, and 3. Predictions using bioinformatics tools suggest that the vaccine is stable, non-allergenic, and provides near total global population coverage (>99%). Because L1 major capsid protein plays a crucial role in HPV infection and its immunogenicity is well documented, it was selected as the source protein in this study. However, the L2 minor capsid protein, inducing low-titer yet cross-neutralizing antibodies, has also been proposed as an alternative to develop broader-spectrum HPV vaccines, and may be a point worth investigating in the future [61].

To ensure high levels of recombinant expression in prokaryotic systems, the GenScript tool was used to confirm the full-length polypeptide vaccine’s suitability for expression in *E. coli* host [13]. The vaccine’s decent solubility propensity score also indicates less accumulation of proteins in the inclusion bodies. With the addition of restriction enzymes and a N-terminal his-tag, the DNA sequence encoding the vaccine was prepared for further studies, which consist of transforming the recombinant plasmid into *E. coli* and performing subsequent tests for immunogenicity [62].

By identifying immunodominant epitopes, this study provides the basis for the development of a peptide-based wart vaccine. However, important questions about enhancement and delivery must be settled sooner or later in the vaccine engineering process. It is known that subunit peptide vaccines can have limited immunogenicity. As such, the incorporation of immunostimulatory adjuvants has often been utilized to enhance vaccine responses [21]. Choosing an ideal adjuvant can help improve innate and acquired immunity, so careful selections should be made based on the host, pathogen, vaccine antigen, and the route of immunization [63]. The arrangement of epitopes and adjuvants should also be subjected to *in silico* optimization [64]. Appropriate consideration of these factors are needed, should the findings in this study be built upon in the future.

## Data Availability Statement

The datasets generated for this study are available on request to the corresponding author.

## Author Contributions

WHT and WHC conceptualized the study. WHT performed the experiments. WHT, CWW, and WHC analyzed the data. WHC and CWW supervised the study. WHT wrote the first draft of the manuscript. All authors contributed to the article and approved the submitted version.

